# Gut microbial community in proboscis monkeys (*Nasalis larvatus*): implications for effects of geographical and social factors

**DOI:** 10.1101/2023.03.14.532648

**Authors:** Lilian Jose, Wanyi Lee, Goro Hanya, Augustine Tuuga, Benoit Goossens, Joseph Tangah, Ikki Matsuda, Vijay Subbiah Kumar

## Abstract

Recent technological advances have enabled comprehensive analyses of the previously uncharacterized microbial community in the gastrointestinal tracts of numerous animal species; however, the gut microbiota of several species, such as the endangered proboscis monkey (*Nasalis larvatus*) examined in this study, remains poorly understood. Our study sought to establish the first comprehensive data on the gut microbiota of free-ranging foregut-fermenting proboscis monkeys and to determine how their microbiota are affected locally by environmental factor, i.e. geographical distance, and social factor, i.e. number of adult females within harem groups and number of adults and subadults within non-harem groups, in a riparian forest in Sabah, Malaysian Borneo. Using 16S rRNA gene sequencing of 264 faecal samples collected from free-ranging proboscis monkeys, we demonstrated the trend that their microbial community composition is not particularly distinctive compared to other foregut and hindgut fermenting primates. The microbial alpha diversity was higher in larger groups and individuals inhabiting diverse vegetation (i.e. presumed to have a diverse diet). For microbial beta diversity, some measures were significant, showing higher values with larger geographical distances between samples. These results suggest that social factors such as increased interindividual interactions, which can occur with larger groups, as well as physical distances between individuals or differences in dietary patterns, may affect the gut microbial communities.

## 1. Introduction

The gastrointestinal tracts of animals are colonized by many microorganisms, forming complex microbial ecosystems [1–3]. In general, animal-associated microbial ecosystems have been reported to have a direct effect on host health, contributing not only to daily energy acquisition through the production of vitamins and short-chain fatty acids but also to the host’s immune system and resistance to pathogens [4–6]. Understanding how the microbial community, which serves important functions in animals, is formed in the gastrointestinal tracts would shed light on the survival strategies of diverse animal species. Historically, the study of the human gastrointestinal microbial community has been extensive [7, 8]. However, with recent advancements in sequencing technology, the ability to analyze gastrointestinal microbial diversity and community structure based on large amplicon libraries of 16S ribosomal RNA (rRNA) genes, primarily using faecal DNA, has prompted additional research in a variety of nonhuman primates [9–12]. With the recent accumulation of such research findings in nonhuman primates, it is becoming increasingly evident that their living environment influences and shapes the gastrointestinal microbial community [13]. For example, within the same primate species, a correlation has been observed between the diversity of dietary items and the microbial community [14–17]. Alternately, as in primate species that live in groups, it has been hypothesized that social factors may influence the establishment of the microbial community, as the horizontal transmission of the microbiota may occur via direct social interactions between individuals within the group or even indirect interaction via shared environments [18–21].

Although the gastrointestinal microbial community, particularly the gut microbiota from faecal samples, has been progressively studied in various primate species over the last decade[10, 22–25], there are still numerous species in the wild for which even the most fundamental microbial community has not been studied. The foregut-fermenting proboscis monkey (*Nasalis larvatus*), a large, sexually dimorphic, arboreal primate [26], is one of these species for which the hindgut microbiota has not been studied in the wild (but see in captivity [27]), though comprehensive analyses of their foregut microbiota have been reported in the wild [15]. As both the foregut and hindgut have been reported to facilitate the digestive fermentation of dietary fibre in the closely related foregut-fermenting primate species, i.e. *Rhinopithecus roxellana* [23], a comprehensive analysis of hindgut bacteria based on faecal samples in proboscis monkeys is essential for fully understanding their digestive physiology.

Proboscis monkeys inhabit various riparian and coastal forest environments, including riverine, mangrove, and peat swamp forests [28], and their dietary patterns have been reported to be flexibly adapted to these environments; diverse diets in riverine forests with higher plant diversity, and low dietary diversity in mangrove forests where plant diversity is extremely low [29–31]. Due to its adaptable feeding habits, the proboscis monkey is an ideal study species for determining how habitat-specific dietary differences influence gut microbiota. The basic component of social structure in proboscis monkeys is as polygynous single male societies (harem group) that assembles with each other in the trees along rivers [32–35], though all-male groups and, occasionally, groups containing more than one male with multiple females are also found [36]. Consequently, it is a fascinating species with which to investigate the relationship between social factors such as different group types with different sexual composition of members and the intestinal microbial community.

We sought to establish the first comprehensive data on the gut microbiota of free-ranging proboscis monkeys and to determine how these gut microbiota were affected at the local scale by environmental factors (e.g. geographical location) and social factors (e.g. sexes with different life histories and group size). In particular, we hypothesise that the observed richness and diversity of gut microbial communities would vary based on the distance of their living areas from the river mouth in the study site and the number of adult females in harem groups. In addition, we propose that beta diversity measures of gut microbiota would correlate with geographical distance between individuals. Previous research has demonstrated that the gut microbiota is frequently influenced by dietary habits; therefore, we anticipated that there would be differences between the gut microbiota of proboscis monkeys living near the river mouth, where deforestation by oil palm plantations is more pronounced, and those living in the upper river areas (see the Supplementary Material 1), where relatively large areas of forest remain at our study site [29]. Indeed, since increased dietary diversity has been reported to increase the diversity of the gut microbial community [e.g. 15, 37, 38], it would be predicted that disturbed habitats would generally experience a reduction in plant diversity [39] and that the monkeys inhabiting these habitats would experience a reduction in dietary diversity. Furthermore, sex differences in social behavior have been associated with sexual biases in the gut microbiota in primates [e.g. 19, 40, 41], and female-to-female grooming is the predominant form of grooming in proboscis monkeys, but rarely between males and females [42, 43]. Thus, female proboscis monkeys are expected to have more contact with more individuals in the group, and accordingly, alpha diversity and composition of their gut microbiota may be expected to be more diverse and/or higher similarity than males. Lastly, given that several reports in primates have stated that direct inter-individual contact is associated with the transmission of gut microbiota [18, 21], it can be predicted that individuals with closer inter-individual distances would have more similar gut microbiota. Additionally, it is likely that individuals in larger groups would have more opportunities for social interaction with more individuals, and as a result, they may possess a more diverse gut microbiota.

## 2. Methods

### 2.1 Study site and subjects

The study was carried out in a riverine forest along the Menanggul River, a tributary of the Kinabatangan River, Sabah, Malaysian Borneo (118° 30′ E, 5° 30′ N), inhabited by eight species of diurnal primates, including our study species, the proboscis monkey. For more than a decade, this area has been a popular tourist destination that attracts boat tours; as a result, the proboscis monkeys were well-habituated to human observers. The study site, in a 4 km stretch from the mouth of the Menanggul River upstream was home to at least 200 proboscis monkeys, organized into 8–10 harem groups with one adult male, multiple adult females and immatures [35, 44], and various non-harem groups, including all-male groups (including solitary males), as well as other groups with multiple males and multiple females [36, 45].The southern portion of the Menanggul River is dominated by secondary forests, while the northern part has been cleared for oil palm plantations, excluding a protected zone along the river (Supplementary Material 1). Daily temperatures in the area were recorded at approximately 24°C (minimum) and 30°C (maximum), with an average annual rainfall of 2,474 mm [29]. The river levels fluctuate by approximately 1 m daily, with seasonal floods causing an average increase of more than 3 m [46].

### 2.2 Faecal sampling

Proboscis monkeys in the lower Kinabatangan Floodplain typically prefer to sleep along the river [47]. Therefore, we conducted a boat survey in the late afternoon to detect proboscis monkeys and record their group composition with GPS coordinates of their sleeping trees. In the early hours of the following morning, while the monkeys were still sleeping, we revisited their sleeping trees. As proboscis monkeys typically defecate shortly before moving into the forest, we carefully searched the ground near their sleeping trees to collect fresh faeces after they had left the sleeping trees [35]. Several harem groups often stayed in close proximity to the trees along the river, and it was sometimes difficult to determine which group the faeces on the forest floor belonged to, but based on the location of the faeces and the location of the group identified during the boat survey on the previous day, we inferred the group of which the individual that had defecated belonged. We only focused on collecting faecal samples presumed to be from adult individuals [48], and between June 2015 and April 2016, a total of 307 samples were opportunistically collected. However, due to the nature of the faeces collection procedure, it was unclear how many groups in the study area the faeces were collected from. The collection was always carried out immediately after defecation, using a sterilized plastic spoon attached to the sampling tube. The spoons were inserted into fresh faeces, and only a small amount of the interior was removed and stored in 5-ml lysis buffer (0.5% sodium dodecyl sulfate, 100 mM ethylenediaminetetraacetic acid (pH 8.0), 100 mM Tris-HCl (pH 8.0), and 10 mM NaCl [16] at room temperature. To prevent duplicate analyses after genetic profiling, faeces from the same individual and those samples of unknown sex were excluded, and thus, the analysis of gut microbiota was performed with 264 faecal samples, i.e. 187 females and 77 males [35].

### 2.3 DNA purification, 16S ribosomal RNA (rRNA) amplification, and sequencing

After bead-beating using the bead crusher (TAITEC, µT-01, Japan) and centrifuged at 4,200 rpm for 5 minutes, 200 μl of lysis buffer-faecal sample mixture was added with 800 μl of InhibitEX buffer of the QIAamp DNA Stool Mini Kit (Qiagen GmbH, Hilden, Germany). Next, the mixture was centrifuged at room temperature for 1 minute at 13,000 rpm. Then, the lysate was transferred to a new 1.5 ml microcentrifuge tube with 25 μl of proteinase K. This was followed by adding 500 μl of Buffer AL and the manufacturer’s protocols to purify the faecal DNA. Next, the DNA concentration was estimated with a Qubit dsDNA HS Assay Kit and a Qubit fluorometer (Thermo Fisher Scientific). We amplified the V3-V4 region of the 16S rRNA gene using the primer utilized in [49] with slight modification as follows: 16S_V34_F 5′-TCGTCGGCAGCGTCAGATGTGTATAAGAGACAG-Ns-CCTACGGGNGGCWG-3,′ and 16S_V34_R 5′-GTCTCGTGGGCTCGGAGATGTGTATAAGAGACAG-Ns-GACTACHVG GG-3′. Subsequently, 3Ns, 4Ns, 5Ns, and 6Ns were inserted in each primer between the specific primer and the adapter to cause an artificial frameshift and improve the sequencing quality [48]. The mixture of these primers was used as forward and reverse primer at concentration of 1 μM. KAPA Pure Beads (KAPA Biosystems, Wilmington, MA) were used to purify the polymerase chain reaction amplicons. The Illumina Nextera XT Index Kit (Illumina, Inc., San Diego, CA) was then used to attach specific dual indices and sequencing adapters to the amplicons for each sample. The resulting products were then combined in equal DNA concentrations to form a pooled sequencing library. Subsequently, the size distribution of the library was then estimated using an Agilent 2100 Bioanalyzer (Agilent Technologies, Inc., La Jolla, CA). The library was then diluted to a concentration of 15 pM and sequenced with a 15% PhiX spike-in on an Illumina MiSeq sequencing platform using the MiSeq Reagent Kit v3 (600 cycles) (Illumina, Inc., San Diego, CA). The resulting read lengths were 301 bp (forward sequences), 8 bp (forward indices), 8 bp (reverse indices), and 301 bp (reverse sequences).

### 2.4 Data analysis

#### Amplicon sequence variants (ASVs) picking and taxonomic identification

The demultiplexed sequences were processed using QIIME2 software [50]. ASVs were generated using the DADA2 pipeline in this software through the dada2 plugin. In this step, the forward and reverse reads were merged. The unmerged reads were also discarded, and chimera were removed. The parameters used were to exclude demultiplexed sequences with quality score below 30 from the downstream analysis to ensure high quality data. The spurious ASVs were also removed by the QIIME2 dada2 plugin. The ASVs were assigned through the ribosomal database project classifier with GreenGenes v13_8 as the reference database for taxonomic identification. Additionally, we used the built-in function align-to-tree-might-fast tree of QIIME2 to construct a phylogenetic tree of the ASVs.

#### Statistical analysis

Statistical analyses were performed in R Version 4.1.1 [51], with the significance level set at 0.05 using the unrarefied dataset with the singletons being removed. Results were reported as means with standard deviation. Alpha and beta diversity was calculated using the R package *phyloseq* [52]. The Kruskal–Wallis test from the R package *dunn.test* was used to analyze the differences in alpha diversity between sexes and group types. Subsequently, we investigated the effects of social and geographical factors on the microbial alpha diversity of individual samples using a linear model. The alpha diversity of each sample was treated as a normally distributed response variable, while the social factor, i.e. the number of adult females for harem groups and the number of adults and subadults for non-harem groups, and the geographical factor, i.e. the location of faecal samples collected along the river, which represented as the distance from the river mouth, were treated as explanatory variables. For all models, we verified that the variance inflation factors were smaller than the cutoff value, i.e. less than 10 [53]. Therefore, the collinearity between independent factors (explanatory variables) did not affect the results. For the model selection, the possible combinations of the explanatory variables were examined and ranked using the Akaike information criterion (AIC) from the *MuMIn* package [54]. Following the published guidelines for wildlife research, we generally selected the best-supported models as those with a *Δ*AIC score of less than 2, where *Δ*AIC = AIC − minimum AIC within the candidate models [55], but as suggested by Burnham et al. [56] which the model with *Δ*AIC of range from 2 to 7 should not be dismissed, we discussed models that were in those ranges as well.

In our multivariate analysis of microbiota composition, we calculated Bray–Curtis dissimilarity, along with weighted and unweighted UniFrac indices with the R package *vegan*. We conducted the permutational multivariate analysis of variance (PERMANOVA, adonis2 function in the vegan package, R software, version 4.1.1) tests to estimate the differences in term of beta diversity. We constructed nonmetric multidimensional scaling (NMDS) for visualization using Bray–Curtis dissimilarity and principal coordinate analysis (PCoA) plots by weighted and unweighted UniFrac indices with the plot_ordination function from package ggplot2. In order to further assess the effect of social and geographical factors and to test their correlation with microbial beta diversity, Mantel tests were conducted by using the mantel function. This function takes three parameters, where the first one is the first distance matrix, followed by the second distance matrix, and the last one is ‘spearman’ which indicates the method that we used to compute the correlation between these two distance matrices. The social factor, known as ‘demographic distance,’ was calculated based on the composition of the differences in the group and the number of individuals within the groups, while the geographical factor, known as ‘geographical distances,’ was calculated based on the distance between different faecal samples (or individuals). In other words, with respect to these two distances, values are higher with more differences in pattern of group composition within a group (i.e. differences in the number of males and females) and with greater distances between different faecal samples (or individuals).

## 3. Results

### 3.1 Phylogenetic profile of the faecal microbiota

In 264 faecal samples, we detected 16,530 ASV, classified into 29 phyla, 67 classes, 95 orders, 116 families, 134 genera, and 50 species. The average number of sequences resulted per sample before filtering through DADA2 pipeline was 124,898, with a maximum of 287,114 and a minimum of 37,712. After filtering, the average was 100,900, with a maximum of 257,018 and a minimum of 10,234.

Figure 1 depicts the general distribution of the top five taxa at the phylum, family, and genus levels. At the phylum level, the top five taxa were consistent across sexes; Bacillota dominated the gut microbial community (male: 81.9%; female: 82.3%), followed by Bacteroidota (male: 8.3%; female: 8.0%), Cyanobacteria (male: 1.5%; female: 1.8%), Pseudomonadota (male: 1.5%; female: 1.3%), and Actinomycetota (male: 1.1%; female: 1.1%) (see Supplementary Material 2 for details). At the family and genus level, the top five patterns of the gut microbial community in both sexes were also consistent (see Supplementary Material 3-4 for details), i.e. family level: Ruminococcaceae (male: 46.3%; female: 44.6%), Lachnospiraceae (male: 14.7%; female: 15.3%), S24-7 (male: 6.8%; female: 6.5%), Christensenellaceae (male: 2.0%; female: 2.2%), and [Mogibacteriaceae] (male: 1.9%; female: 1.8%); genus level: *Oscillospira* (male: 10.4%; female: 10.2%), *Ruminococcus* (male: 4.7%; female: 4.5%), *Dorea* (male: 2.3%; female: 2.1%), and *Blautia* (male: 1.2%; female: 1.4%). In addition, we found that the pattern of the top five taxa (at the phylum, family, and genus levels) was consistent across different group types (harem and non-harem groups) (see Supplementary Material 5-7). Finally, despite the information available on annotated bacterial taxa at the phylum, family and genus level, it would be worth noting the proportion of unassigned taxa, which were 3.18–3.38%, 16.02–17.14%, and 71.82– 72.62%, respectively.

**Figure 1.**
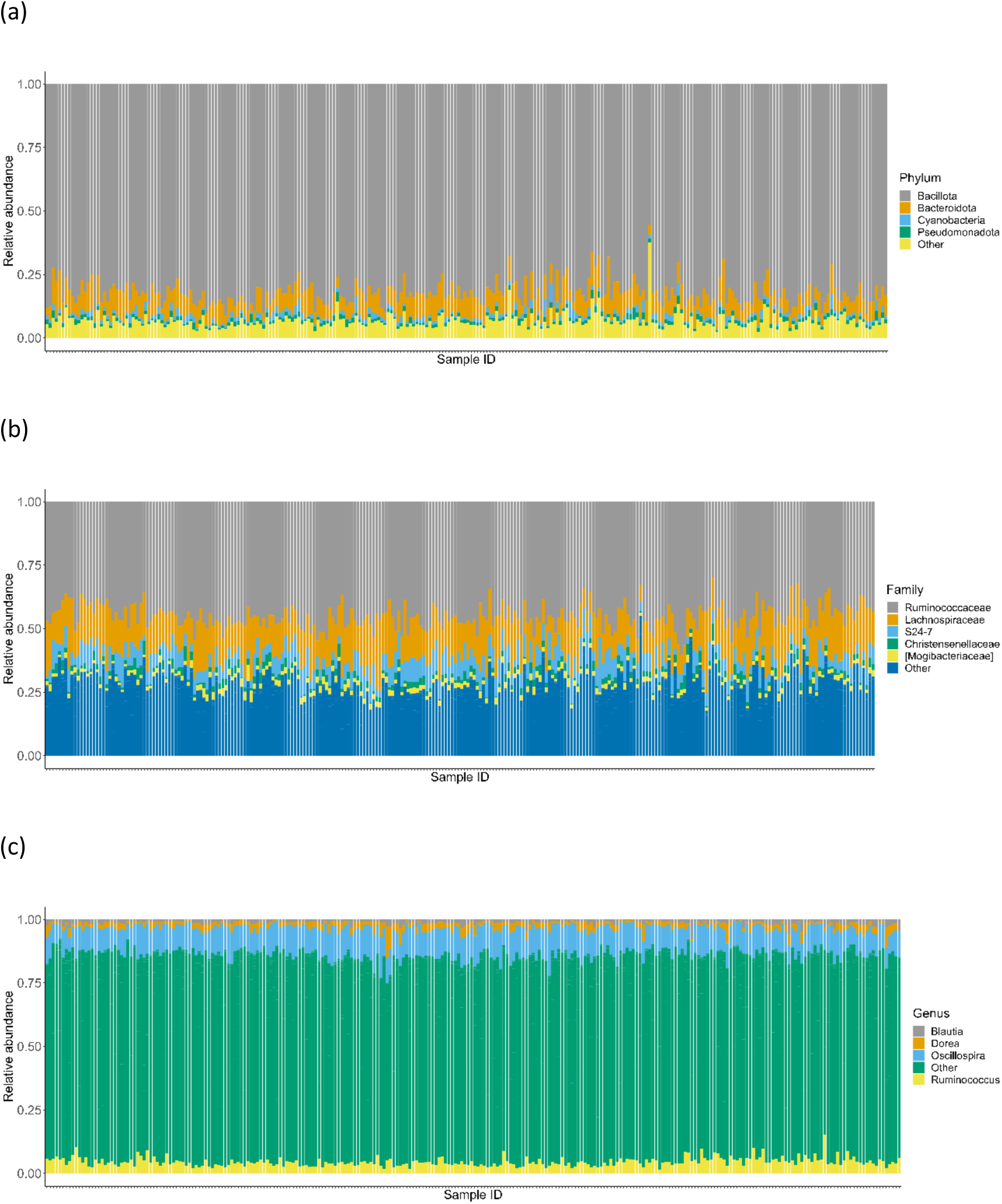
The general pattern of the top five taxa. These were analyzed at the phylum (a), family (b), and genus (c) levels.

### 3.2 Gut microbial diversity

#### Alpha diversity

The mean observed richness and Shannon diversity index (*H’*) of the gut microbiota in all faecal samples were 400.6 ± 51.5 and 5.0 ± 0.18, respectively. There were no significant differences in observed richness between sexes (Kruskal–Wallis χ² = 2.77, df = 1, *p = 0*.10; Figure 2a), nor between the group types (harem and non-harem groups) (Kruskal–Wallis χ² = 2.97, df = 1, *p = 0*.08; Figure 2a). Shannon diversity index did not differ significantly by sex (Kruskal–Wallis χ² = 1.81, df = 1, *p = 0*.18; Figure 2b) or group type (Kruskal–Wallis χ² = 0.25, df = 1, *p = 0*.62). Richness values ranged between 161 and 661. In contrast, the Shannon diversity index values ranged between 4.17 and 5.48.

**Figure 2.**
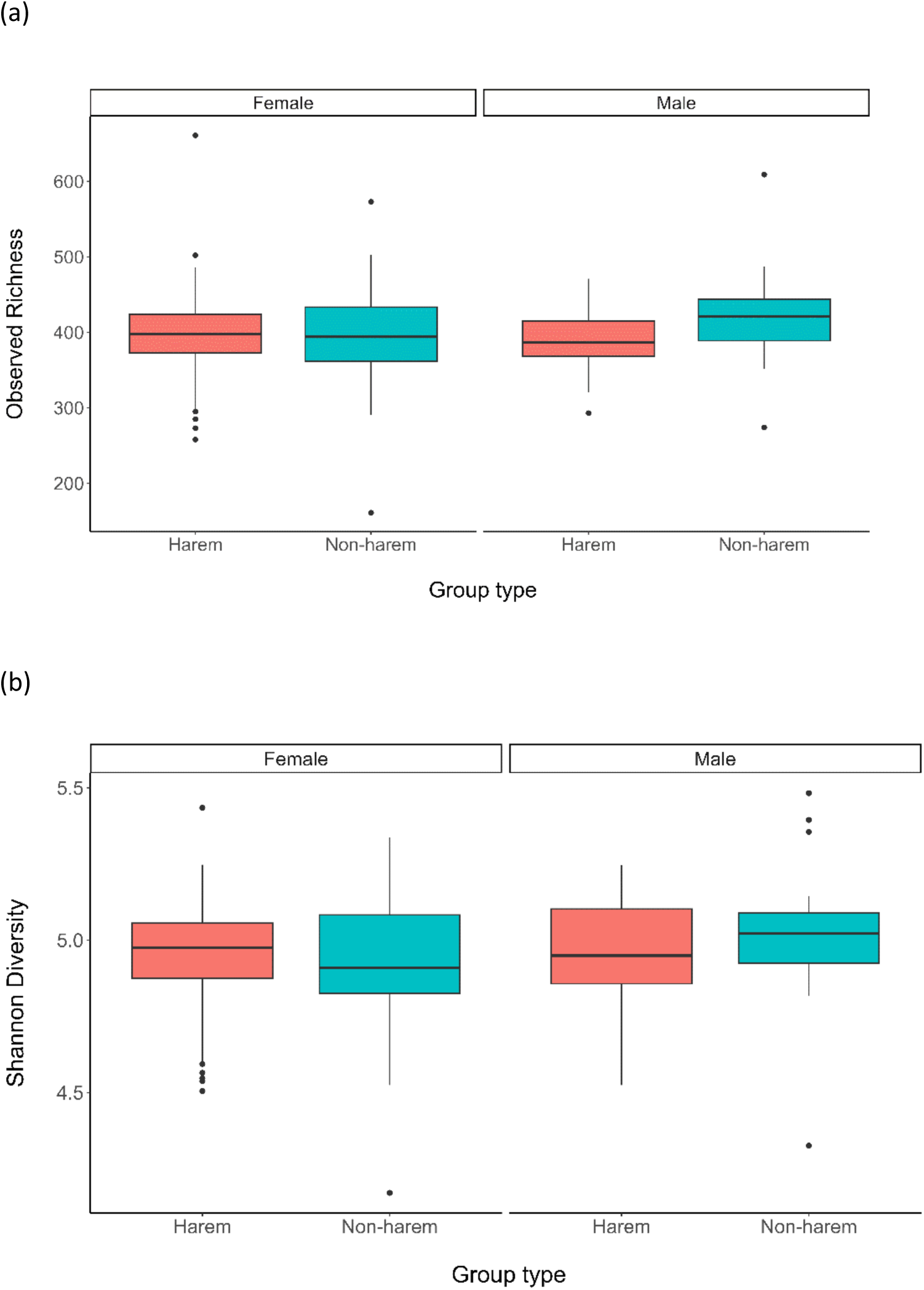
Comparison of the mean observed richness (a) and the Shannon diversity index (b) for different sexes and different groups. The central line of the box represents the median, and the lower and upper bounds of the box represent the first and third quartiles.

The best-fit model to explain the observed richness, as determined by AIC, included both the number of adult females in the harem groups and the location of the collected samples represented as the geographical distance from the mouth of the river (Table 1 and Figures 3a), although the *Δ*AIC value of the following model, which included only the number of adult females, was also <7.0. The observed richness increased as the number of adult females in the harem groups, and the distance from the river mouth increased (upper river). The linear models for Shannon diversity index revealed a similar pattern to the observed richness in harem groups, with a positive effect on the number of adult females and geographical distance (Table 1 and Figures 3b).

**Figure 3.**
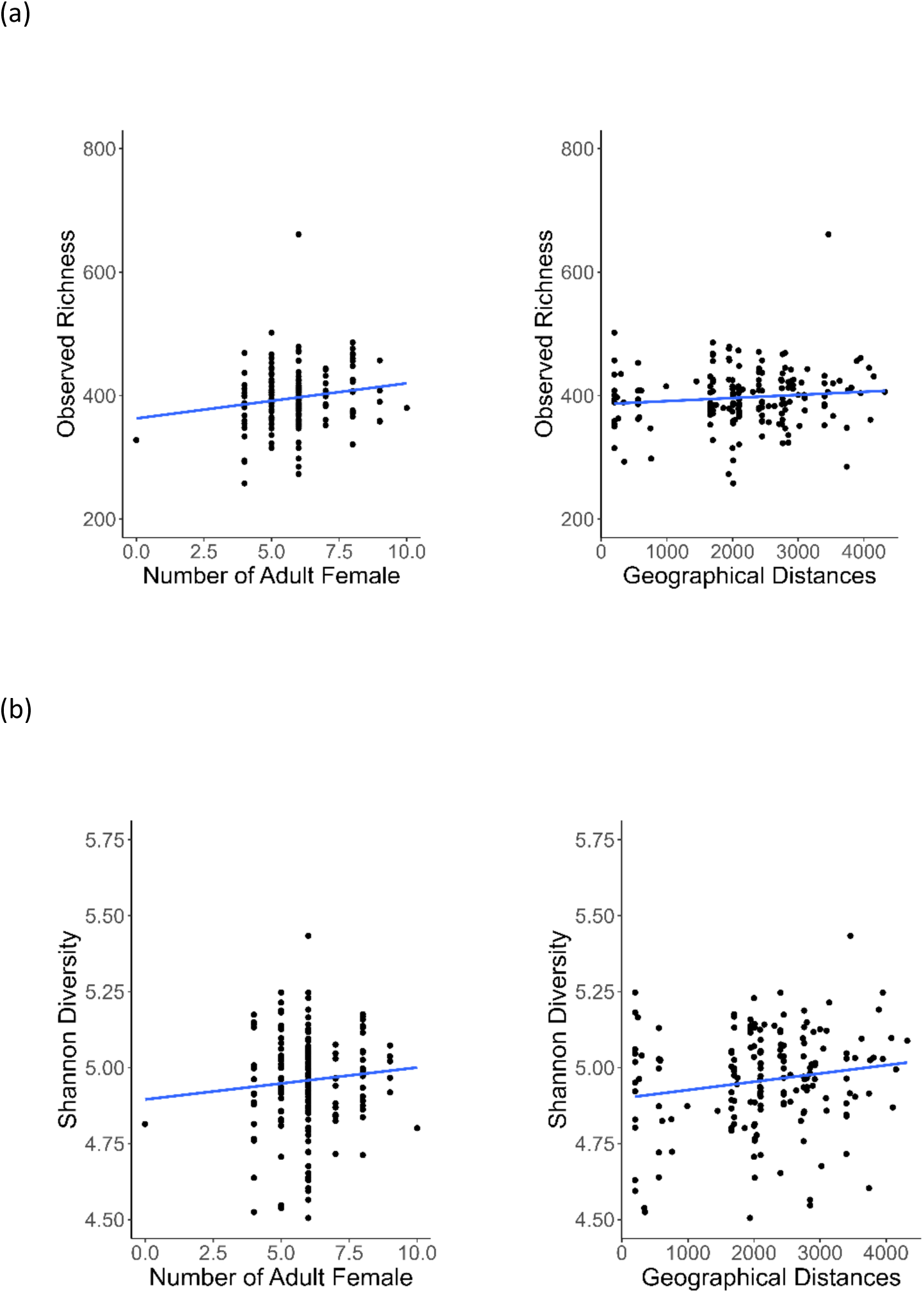
Relationship of alpha diversity index, i.e. observed richness (a) and Shannon diversity index (b), with number of adult females within harem groups and location of faecal samples collected representing as geographical distance from the river mouth, based on the model selection Table 1.

**Table 1.** Summary of model selection for harem groups using linear models. This method was used to investigate whether observed richness (A) and Shannon diversity index (B) were affected by social factor, i.e. the number of adult females within harem groups, and geographical factor, i.e. the location of faecal samples collected along the river, which represented as the distance from the river mouth.

| (A) | Intercept | Adult female | Distance from river mouth | df | Log-likelihood | AIC | Δ AIC | Weight |
| --- | --- | --- | --- | --- | --- | --- | --- | --- |
|  | 351.5 | 5.758 | 0.005165 | 4 | -902.163 | 1812.3 | 0.00 | 0.408 |
|  | 363.0 | 5.691 |  | 3 | -903.226 | 1812.5 | 0.13 | 0.383 |
|  | 396.9 |  |  | 2 | -905.517 | 1815.0 | 2.71 | 0.105 |
|  | 386.1 |  | 0.005031 | 3 | -904.535 | 1815.1 | 2.74 | 0.103 |
| **(B)** |  |  |  |  |  |  |  |  |
|  | 4.899 |  | 2.721e-05 | 3 | 72.000 | -138.0 | 0.00 | 0.463 |
|  | 4.834 | 0.01087 | 2.746e-05 | 4 | 72.715 | -137.4 | 0.57 | 0.348 |
|  | 4.958 |  |  | 2 | 69.568 | -135.1 | 2.86 | 0.111 |
|  | 4.895 | 0.01052 |  | 3 | 70.219 | -134.4 | 3.56 | 0.078 |

Within the non-harem groups, however, model selection of the model for the investigation of whether the observed richness and Shannon diversity index were affected by social and geographical factors, i.e. the number of individuals, including adult males and females and subadult males, and the location of the collected samples, represented as the geographical distance from the river mouth, indicated that the null model was the best (Table 2). The second- and third-best models, with observed richness and Shannon diversity index, incorporated social and geographical factors with *Δ*AIC values below the cutoff of 7.0. This result indicates that a similar pattern was generally observed in non-harem groups, albeit with weaker effects than in harem groups.

**Table 2.** Summary of model selection for non-harem groups using linear models. This method was used to investigate whether observed richness (A) and Shannon diversity index (B) were affected by social factor, i.e. the number of individuals, including adult males and females and subadult males within groups, and geographical factor, i.e. the location of faecal samples collected along the river, which represented the distance from the river mouth.

| (A) | Intercept | Number of adults and subadults | Distance from river mouth | df | Log-likelihood | AIC | $\Delta$ AIC | Weight |
| --- | --- | --- | --- | --- | --- | --- | --- | --- |
|  | 407.6 |  |  | 2 | -504.772 | 1013.5 | 0.00 | 0.525 |
|  | 405.2 |  | 0.001834 | 3 | -504.716 | 1015.4 | 1.89 | 0.204 |
|  | 401.7 | 1.185 |  | 3 | -504.763 | 1015.5 | 1.98 | 0.195 |
|  | 396.7 | 1.678 | 0.001987 | 4 | -504.699 | 1017.4 | 3.85 | 0.076 |
| **(B)** |  |  |  |  |  |  |  |  |
|  | 4.969 |  |  | 2 | 18.998 | -34.0 | 0.00 | 0.503 |
|  | 4.954 |  | 1.146e-05 | 3 | 19.190 | -32.4 | 1.62 | 0.224 |
|  | 4.946 | 0.004727 |  | 3 | 19.010 | -32.0 | 1.98 | 0.187 |
|  | 4.915 | 0.007743 | 1.217e-05 | 4 | 19.222 | -30.4 | 3.55 | 0.085 |

#### Beta diversity

According to PCoA using weighted and unweighted UniFrac and NMDS plots with Bray-Curtis dissimilarity, individuals of different sexes or group types did not exhibit visually distinguishable patterns (Figure 4). Conversely, PERMANOVA analysis revealed weak but significant differences between both sexes (PERMANOVA, Bray–Curtis, R² = 0.0049, *p =* 0.021; unweighted UniFrac, R² = 0.0045, *p =* 0.009) and between group types (PERMANOVA, Bray–Curtis, R² = 0.0053, *p =*0.004; unweighted UniFrac, R² = 0.0045, *p =* 0.01) on the gut microbiota.

**Figure 4.**
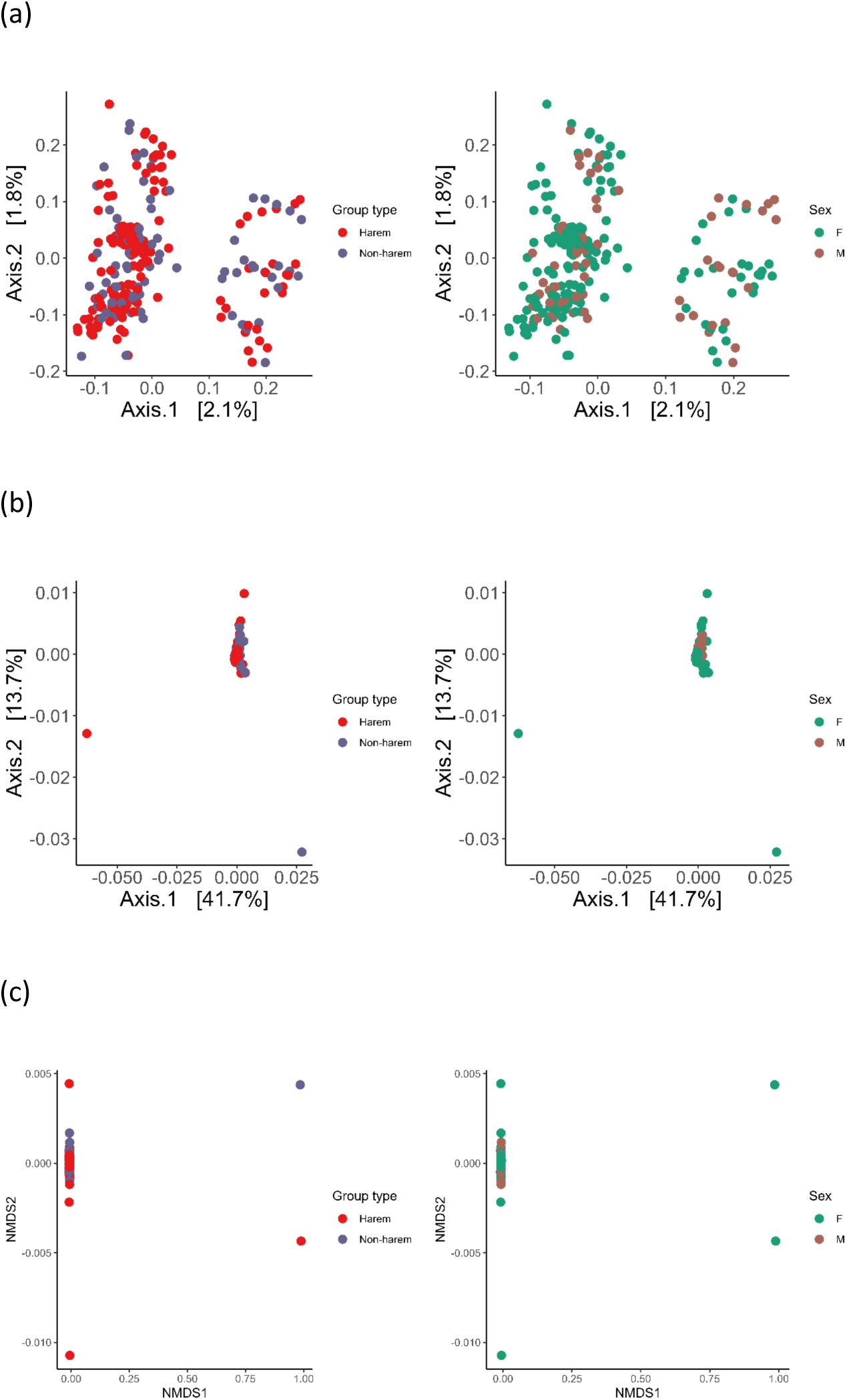
Unweighted UniFrac (a) and weighted UniFrac (b) principal coordinate analysis (PCoA) plots and Bray–Curtis nonmetric multidimensional scaling (NMDS) plot (c), analyzed by group type, i.e. harem and non-harem, and sex, i.e. F: female; M: male.

In order to examine the correlation between beta diversity measures and the two factors, i.e. geographical distances and social factors, Mantel test was conducted separately on a dataset containing only samples from harem groups and one containing only samples from non-harem groups. Mantel tests performed with different beta diversity measures revealed significant positive correlations for the harem group between weighted UniFrac and geographical distances from different individuals (r = 0.09, *p* = 0.01) but not between unweighted UniFrac and geographical distances from different individuals (r = 0.02, *p* = 0.21). This indicated that individuals living further apart had a gut microbial community that was less similar, although the explanatory power of geographic distance was relatively weak. In contrast, Bray–Curtis dissimilarity measures of diversity, which do not account for the phylogeny of microbial composition, did not exhibit a significant correlation with geographic distance (r = 0.05, *p* = 0.09). In addition, there was no correlation between the different beta diversity measures and the distance of group composition between the various harem groups (weighted UniFrac: r = 0.10, *p* = 0.09; unweighted UniFrac: r = 0.02, *p* = 0.23; Bray–Curtis: r = 0.002, *p* = 0.48).

For samples from non-harem groups, weighted UniFrac (r = 0.13, *p* = 0.02) and Bray– Curtis dissimilarity (r = 0.23, *p* = 0.001) were significantly correlated with geographical distance, whereas there was no significant correlation between unweighted UniFrac and geographical distance (r = 0.03, *p* = 0.19). All the beta dissimilarity indices exhibited a significant correlation with the similarity distance of the group composition between different groups (weighted Unifrac: r = 0.17, p = 0.03; unweighted Unifrac: r = 0.10, *p* = 0.04; Bray-Curtis: r = 0.13, *p* = 0.04).

## 4. Discussion

### 4.1 Overall gut microbial composition

The present study was the first comprehensive analysis of the intestinal microbial community based on faeces from more than 250 free-ranging proboscis monkeys. At the phylum level, the composition of their gut microbial community was nearly identical to that of other foregut-fermenting primates. Bacillota and Bacteroidota were predominant in *Colobus polykomos*, *Procolobus badius*, *P. verus*, *Rhinopithecus brelichi,* and *R. roxellana* [12, 57–59], with the exception of *Pygathrix nemaeus*, which includes Bacteroidota in the top five, but Bacillota and Mycoplasmatota dominate the top two phylum [38]. This trend is not exclusive to foregut-fermenters but is also observed in hindgut-fermenters, such as *Cercocebus atys*, *Cercopithecus campbelli*, *C. diana*, *C. petaurista*, *Chlorocebus sabaeus*, *Lemur catta*, *Macaca fuscata*, *M. mulatta*, *M. thibetana*, *Pan troglodytes*, *Propithecus verreauxi* and *Theropithecus gelada* [12, 57, 60–64].

At the family level, the predominance of Ruminococcaceae, or Lachnospiraceae is comparable to that of the foregut-fermenting *R. brelichi* and *R. roxellanae*. *Macaca fuscata*, a hindgut-fermenting primate, exhibited a similar trend [60], and Ruminococcaceae were frequently the most dominant in other hindgut-fermenting primates [12, 61, 62]. Although the proportion taxonomically assigned at the genus level was limited, the top two genera, *Oscillospira* and *Ruminococcus*, are listed among the top five in *P. nemaeus*, a foregut-fermenter as well as in the proboscis monkey. *Ruminococcus,* in particular, is a common genus, as it frequently ranks first not only in foregut-fermenters but also in hindgut-fermenters [60, 62]. However, *Blautia* is not among the top genera in *P. nemaeus*, but it is among the top genera [60, 62] in the hindgut fermenters. In light of these findings, the composition of the gut microbial community of proboscis monkeys would not be significantly different from that of other foregut-/hindgut-fermenting primates.

In the light of these findings, the composition of the gut microbial community of the proboscis monkey is unlikely to differ significantly from that of other foregut and hindgut fermenting primates. However, when the microbiota is subdivided from the phylum level to the genus level, the degree of overlap obviously decreases, and species-specific microbial composition and bacterial species are found probably due to more species-specific digestive physiology, dietary patterns related to living environment and/or, sociality of host animals, e.g. lactic acid bacteria, *Lactobacillus nasalidis*, specific only to the proboscis monkey [65, 66].

### 4.2 Microbial patterns in relation to social factors

There were no differences in the alpha diversity index (number of ASV and Shannon diversity index) of the proboscis monkey gut microbiota between sexes or group types, indicating that alpha diversity is not affected by differences in social factors such as sex differences in life history and/or social composition of individuals. Conversely, significant differences in beta diversity were observed between the sexes, suggesting that differences in life history and the frequency of social interactions may have influenced the composition of the gut microbiota. Significant differences in beta diversity between group types may also reflect differences of life history in the sexes; the basis for the differences in gut microbiota between group types with different male and female compositions within a group may be related to the differences in such life histories. At maturity, females transfer from their natal harem groups to other harem groups, whereas males disperse from their natal harem groups in the early stages to join all-male groups [32, 42, 67, 68]. Additionally, grooming within harem groups occurs primarily between females, with males rarely participating [43, 69]. Consequently, it is likely that these differences in the life histories of the sexes and in the frequency of social interactions between the sexes influence the composition of the gut microbiota in proboscis monkeys. Several other nonhuman primates have been reported to exhibit sexual biases in the gut microbiota, which could be attributed to differences in such interactions and life histories between sexes as a result of group living, e.g. *Callithrix jacchus* [40], *Rhinopithecus bieti* [41], and *Alouatta pigra* [19], though such sexual differences are little in some species, e.g. *Propithecus verreauxi* [63]. In contrast, it is difficult to determine its ecological significance without knowing how differences in microbiota composition affect factors such as food digestion and immunity. As expected, the positive correlation between the number of females in harem groups and the alpha diversity index suggests that increased individual interactions result in an increase in alpha diversity. This result is consistent with previous research indicating that direct physical contact between social partners is a major factor in the transmission of gut microbiota [19, 21, 40].

### 4.3 Microbial patterns in relation to geographical factors

In addition to the number of females within the harem groups, individuals who resided in areas further upstream of the river mouth at our study site had higher alpha diversity indices. This may be due to differences in the food diversity consumed by proboscis monkeys in upstream and downstream regions. In general, individuals with a more varied diet have a greater variety of symbiotic bacteria in their gastrointestinal tracks [15, 37, 38]. Indeed, given that the downstream area of the study site is more heavily affected by deforestation [29] and has often lower plant diversity with lower potential food sources for proboscis monkeys, especially on the north side of the river (Supplementary Material 9), individuals in the upstream area may have had access to a greater variety of food sources, resulting in a tendency for a higher alpha diversity index in their gut microbiota. Nevertheless, to strengthen this conclusion, differences in vegetation between upstream and downstream areas would require a more quantitative investigation and comparison with the degree of gut microbial diversity obtained in the present study.

In the non-harem groups, a similar relationship was observed between alpha diversity and the number of individuals in the group and the geographical conditions they inhabited, although this trend was weaker with insignificance. This difference may be attributable to the fact that non-harem groups, particularly all-male groups, exploit a broader range of riparian habitats than harem groups [35], which not only facilitate a more diverse diet but also direct/indirect interactions with conspecifics, coupled with increased such interactions with various organisms in the forest elsewhere, possibly leading to the horizontal transmission of the microbiota [70, 71]. However, not only the social relationships between individuals within non-harem groups but also those with various organisms remain unclear, making further discussion impossible at this time. The key to advancing this discussion in the future would be the collection of additional ecological and social observation data on non-harem groups.

There was a weaker but significant correlation between the physical distance between individuals and their similarity in the composition of their intestinal microbial communities, regardless of the group type. It should be noted, however, that the significance of the correlation varied slightly between the various group types based on the various indices, namely weighted UniFrac, unweighted UniFrac, and Bray–Curtis. As previously mentioned, differences in vegetation between upstream and downstream areas of this study site may influence the alpha diversity of individual proboscis monkeys; it cannot be ruled out that the physical distance between individuals may lead to differences in diets consumed, which may, in turn, lead to differences in the composition of the intestinal microbial community between individuals. However, no correlation was found between similarities in group composition and similarities in gut microbial community composition. Compared to seasonal and clumped food sources such as fruits and flowers, the higher ubiquitous and abundant availability of leaves as the main food source for proboscis monkeys in this study site [31, 72] and the lack of a clear hierarchy between individuals, which occurs particularly within harem groups [42]. This may have contributed to the lower likelihood of dietary bias across group composition in proboscis monkeys, and hence there might be no differences in the similarity of the composition of gut microbial community between individuals. Conversely, there have been no studies on the hierarchy among males in non-harem groups, particularly in all-male groups, and the finding that significant differences were observed in non-harem groups by all the beta dissimilarity indices may suggest that there may be a severe hierarchy among males, which resulted in differences in their dietary composition. To verify this, however, more comprehensive behavioral observations of all-male groups are required.

### 4.4 Outlook

We succeeded not only in determining the general trends in the intestinal microbiota of proboscis monkeys but also in determining how social and geographical factors affected this microbial community. In contrast, while we characterized such differences in gut microbial community at the local level among individuals, we have not been able to elucidate how these differences act as an advantage to the survival of each individual. Future research must examine how differences in the gut microbial community between individuals influence their feeding strategy through more detailed functional analysis, such as by isolating and cultivating characteristic strains [66]. Furthermore, considering that the gut microbiota also changes with diet in primates [e.g. 60, 62, 73, 74], there remains a need to compare the dietary patterns and gut microbiota of the individuals in a real-time manner, which we could not demonstrate in the current study, which was a limitation. When analyzing gut microbiota from faeces, only a portion of the faeces is utilized for that analysis, and therefore many other portions are discarded without being utilized. However, with recent metabarcoding techniques [75–77], it may be possible to estimate dietary patterns from that remaining faecal sample. Hence, the analysis of inter-individual differences in gut microbial community related to the different dietary patterns of each individual would be relatively feasible in the future study. Lastly, the present study suggested that the degree of deforestation may influence the gut microbiota, although quantitative comparisons of these effects were not possible. While traditional vegetation surveys are often beneficial to understand the relationship between environment and animal ecology [39, 78–80], advanced technologies such as the recently developing LiDAR technology [81] may allow more extensive and comprehensive habitat assessments which, when coupled with analysis of gut microbial data of target animals as in the present study, may reveal more reliable interactions between their living environment and gut microbiota. In sum, with advancements in technology for the analysis of gut microbiota, it may be possible in the future to transition from non-invasively collected free-ranging primate faeces to studies that contribute to animal conservation, such as assessing the effects of forest disturbance. Multiple studies have demonstrated their potential use in animal conservation [15, 17, 37, 82, 83].

#### Ethics

This study was reviewed and approved by the Sabah Biodiversity Council (Access License Reference: JKM/MBS.1000-2/2 JLD.5) and was conducted in compliance with animal care regulations applicable to Malaysian laws. This study is reported in accordance to ARRIVE guidelines (https://arriveguidelines.org).

#### Data accessibility

The original contributions presented in the study are included in the article/Supplementary Material, further inquiries can be directed to the corresponding author/s. The sequencing data have been deposited to NCBI under <u>PRJNA928786</u>.

#### Declaration of AI use

We have not used AI-assisted technologies in creating this article.

#### Authors’ contributions

B.G., I.M. and V.S.K. conceptualized the initial idea; I.M. collected faecal samples;

A.T. and J.T. arranged the sampling in the field; L.J., G.H., and W.L. conducted a genetic experiment. I.L, W.L., and I.M. performed and interpreted the statistical analyses; L.J., I.M., and V.S.K. drafted the manuscript. All authors contributed to the final version of the manuscript.

#### Funding

This study was partially funded by Japan Science and Technology Agency Core Research for Evolutional Science and Technology 17941861 (#JPMJCR17A4) and Japan Society for the Promotion of Science KAKENHI (#26711027, #15K14605, #19H03308 and 24H00774 to IM; #23H02563 to GH).

## Acknowledgements

We express our sincere thanks to the Sabah Biodiversity Centre and the Sabah Wildlife Department for granting permission to carry out this research. We are also deeply indebted to the Sabah Forestry Department for facilitating the use of their facilities in the field. We are grateful to the researchers at the Biotechnology Research Institute (UMS) and the Inuyama Campus of Kyoto University for the knowledge transfer along the project. LJ and VSK thank Universiti Malaysia Sabah for managing funds grant numbers LPA2001 and LPA2101. IM thanks the research assistants, especially Asnih Binti Etin and Jasrudy Bin Mandu, for their support in the field.

[14–17]

