## Supplementary Materials 1-9 for "Gut microbial community in proboscis monkeys (*Nasalis larvatus*): implications for effects of geographical and social factors"

Jose *et al.* Gut microbial community in proboscis monkeys (*Nasalis larvatus*): implications for effects of geographical and social factors

Supplementary Material 1 Study site in Sabah, Borneo, Malaysia

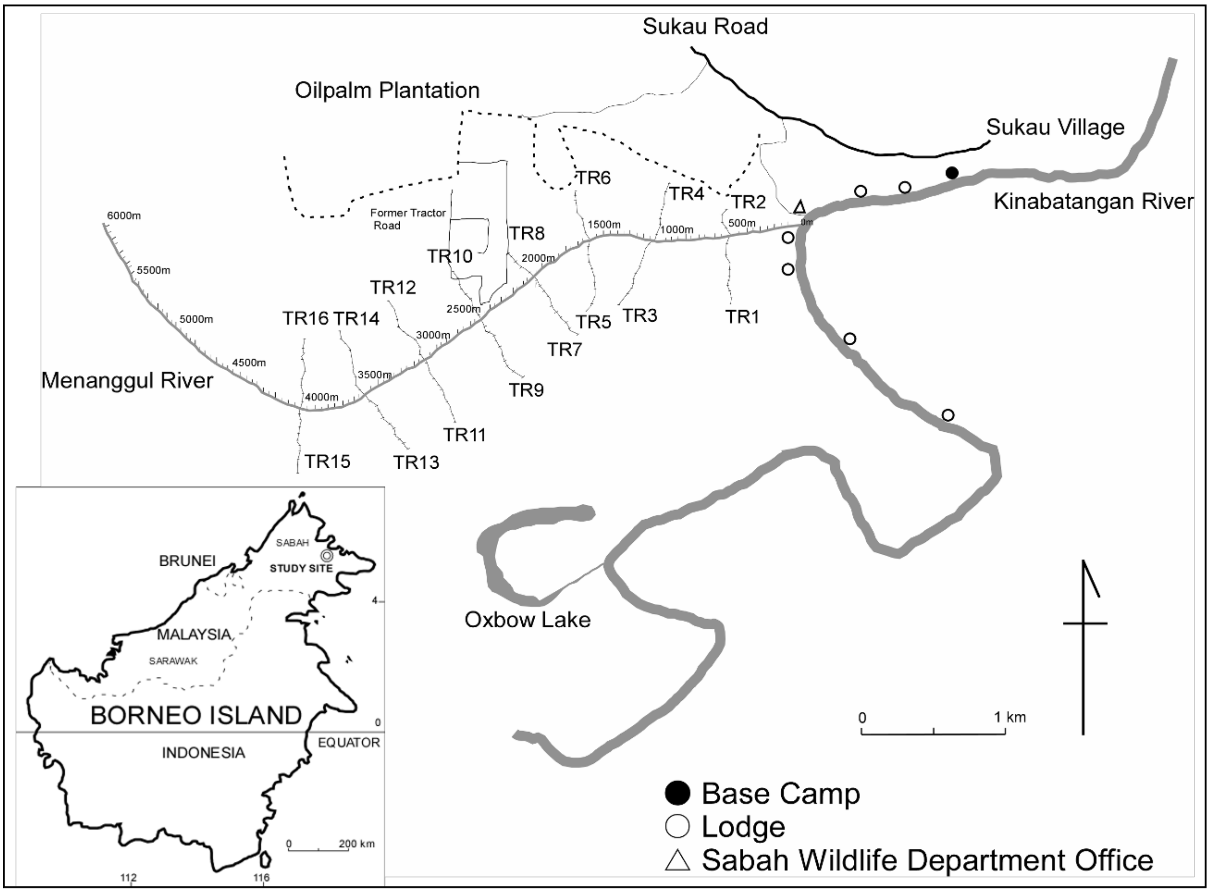


* TR 1 – 16: Trail code

#### Supplementary Materials

Jose *et al.* Gut microbial community in proboscis monkeys (*Nasalis larvatus*): implications for effects of geographical and social factors

Supplementary Material 2: Relative abundance of commonly observed phyla of the faecal microbiome of proboscis monkeys by sex

| **Sex** | | | |
|---|---|---|---|
| **Male** | | **Female** | |
| **Phylum** | **% ± SD** | **Phylum** | **% ± SD** |
| Bacillota | 81.85% ± 0.019 | Bacillota | 82.29% ± 0.019 |
| Bacteroidota | 8.30% ± 0.012 | Bacteroidota | 7.96% ± 0.013 |
| Cyanobacteria | 1.50% ± 0.002 | Cyanobacteria | 1.80% ± 0.006 |
| Pseudomonadota | 1.52% ± 0.002 | Pseudomonadota | 1.33% ± 0.0015 |
| Actinomycetota | 1.09% ± 0.0019 | Actinomycetota | 1.07% ± 0.003 |

#### Supplementary Materials

Jose *et al.* Gut microbial community in proboscis monkeys (*Nasalis larvatus*): implications for effects of geographical and social factors

Supplementary Material 3: Relative abundance of commonly observed family of the faecal microbiome of proboscis monkeys by sex

**Sex**

**Male**

| **Family** | **% ± SD** |
|---|---|
| *Ruminococcaceae* | 46.25% ± 0.0235 |
| *Lachnospiraceae* | 14.67% ± 0.0140 |
| S24-7 | 6.81% ± 0.0357 |
| *Christensenellaceae* | 1.96% ± 0.0162 |
| *[Mogibacteriaceae]* | 1.90% ± 0.0072 |

**Female**

| **Family** | **% ± SD** |
|---|---|
| *Ruminococcaceae* | 44.61% ± 0.0236 |
| *Lachnospiraceae* | 15.34% ± 0.0177 |
| S24-7 | 6.50% ± 0.0385 |
| *Christensenellaceae* | 2.20% ± 0.0135 |
| *[Mogibacteriaceae]* | 1.84% ± 0.0157 |

#### Supplementary Materials

Jose *et al.* Gut microbial community in proboscis monkeys (*Nasalis larvatus*): implications for effects of geographical and social factors

Supplementary Material 4: Relative abundance of commonly observed genera of the faecal microbiome of proboscis monkeys by sex

| Sex | | | | |
|---|---|---|---|---|
| **Male** | | | **Female** | |
| **Genus** | **% ± SD** | | **Genus** | **% ± SD** |
| *Oscillospira* | 10.44% ± 0.0234 | | *Oscillospira* | 10.17% ± 0.0296 |
| *Ruminococcus* | 4.69% ± 0.0227 | | *Ruminococcus* | 4.50% ± 0.0346 |
| *Dorea* | 2.25% ± 0.0189 | | *Dorea* | 2.06% ± 0.0291 |
| *Blautia* | 1.16% ± 0.00978 | | *Blautia* | 1.39% ± 0.0185 |

#### Supplementary Materials

Jose *et al.* Gut microbial community in proboscis monkeys (*Nasalis larvatus*): implications for effects of geographical and social factors

Supplementary Material 5: Relative abundance of commonly observed phyla of the faecal microbiome of proboscis monkeys by group type

| Group Type | | | |
|---|---|---|---|
| Harem | | Non-harem | |
| **Phylum** | **% ± SD** | **Phylum** | **% ± SD** |
| Bacillota | 82.56% ± 0.0188 | Bacillota | 81.43% ± 0.0186 |
| Bacteroidota | 7.83% ± 0.0099 | Bacteroidota | 8.49% ± 0.013 |
| Cyanobacteria | 1.76% ± 0.0059 | Cyanobacteria | 1.60% ± 0.0044 |
| Pseudomonadota | 1.37% ± 0.00248 | Pseudomonadota | 1.42% ± 0.0018 |
| Actinomycetota | 1.06% ± 0.0031 | Actinomycetota | 1.11% ± 0.0025 |

#### Supplementary Materials

Jose *et al.* Gut microbial community in proboscis monkeys (*Nasalis larvatus*): implications for effects of geographical and social factors

Supplementary Material 6: Relative abundance of commonly observed families of the faecal microbiome of proboscis monkeys by group type

| Group Type | | | |
|---|---|---|---|
| **Harem** | | **Non-harem** | |
| **Family** | **% ± SD** | **Family** | **% ± SD** |
| Ruminococcaceae | 45.30% ± 0.0236 | Ruminococcaceae | 44.70% ± 0.0234 |
| Lachnospiraceae | 14.88% ± 0.0176 | Lachnospiraceae | 15.64% ± 0.0148 |
| S24-7 | 6.42% ± 0.0282 | S24-7 | 6.91% ± 0.0382 |
| Christensenellaceae | 2.21% ± 0.0140 | Christensenellaceae | 1.97% ± 0.0160 |
| [Mogibacteriaceae] | 1.86% ± 0.0154 | [Mogibacteriaceae] | 1.86% ± 0.00811 |

#### Supplementary Materials

Jose *et al.* Gut microbial community in proboscis monkeys (*Nasalis larvatus*): implications for effects of geographical and social factors

Supplementary Material 7: Relative abundance of commonly observed genera of the faecal microbiome of proboscis monkeys by group type

| Group Type | | | |
|---|---|---|---|
| **Harem** | | **Non-harem** | |
| **Genus** | **% ± SD** | **Genus** | **% ± SD** |
| *Oscillospira* | 10.32% ± 0.0296 | *Oscillospira* | 10.10% ± 0.0269 |
| *Ruminococcus* | 4.56% ± 0.0254 | *Ruminococcus* | 4.54% ± 0.0254 |
| *Dorea* | 1.93% ± 0.0180 | *Dorea* | 2.45% ± 0.0272 |
| *Blautia* | 1.30% ± 0.0138 | *Blautia* | 1.37% ± 0.0185 |

#### Supplementary Materials

Jose *et al.* Gut microbial community in proboscis monkeys (*Nasalis larvatus*): implications for effects of geographical and social factors

Supplementary Material 8: Information of collected faecal samples

| Sample ID | Group type | Sex | Distance from river mouth (m) | Adult male | Adult female | Subadult male | Subadult female |
|---|---|---|---|---|---|---|---|
| s1 | Non-harem | Female | 3300 | 2 | 1 | 3 | 0 |
| s2 | Harem | Female | 3090 | 1 | 5 | 0 | 0 |
| s3 | Harem | Female | 2800 | 1 | 5 | 0 | 0 |
| s4 | Harem | Female | 2740 | 1 | 5 | 0 | 0 |
| s5 | Harem | Female | 340 | 1 | 5 | 0 | 0 |
| s6 | Harem | Female | 260 | 1 | 5 | 0 | 0 |
| s7 | Harem | Female | 4100 | 1 | 7 | 0 | 0 |
| s8 | Harem | Male | 4080 | 1 | 5 | 0 | 0 |
| s9 | Non-harem | Male | 3560 | 2 | 0 | 3 | 0 |
| s10 | Non-harem | Female | 3000 | 2 | 4 | 0 | 0 |
| s11 | Harem | Female | 2550 | 1 | 6 | 0 | 0 |
| s13 | Harem | Female | 240 | 1 | 6 | 0 | 0 |
| s14 | Harem | Female | 2765 | 1 | 5 | 0 | 0 |
| s15 | Harem | Female | 1740 | 1 | 6 | 0 | 0 |
| s19 | Harem | Male | 3020 | 1 | 6 | 0 | 0 |
| s20 | Harem | Female | 2850 | 1 | 5 | 0 | 0 |
| s22 | *Non-harem | Male | 650 | 1 | 5 | 0 | 0 |
| s25 | Harem | Female | 3050 | 1 | 5 | 0 | 0 |
| s26 | Non-harem | Female | 3000 | 2 | 0 | 3 | 0 |
| s27 | Harem | Female | 2800 | 1 | 6 | 0 | 0 |
| s28 | Harem | Male | 2400 | 1 | 6 | 0 | 0 |
| s29 | Non-harem | Female | 1750 | 2 | 0 | 3 | 0 |
| s30 | Non-harem | Male | 1500 | 2 | 0 | 3 | 0 |
| s31 | Harem | Female | 760 | 1 | 6 | 0 | 0 |
| s32 | Non-harem | Female | 2410 | 2 | 0 | 3 | 0 |
| s33 | Non-harem | Female | 1750 | 0 | 0 | 3 | 0 |
| s34 | Harem | Male | 1700 | 0 | 0 | 0 | 0 |
| s35 | Harem | Female | 750 | 1 | 5 | 0 | 0 |
| s36 | Harem | Male | 350 | 1 | 4 | 0 | 0 |
| s39 | Harem | Female | 3740 | 1 | 6 | 0 | 0 |
| s40 | Harem | Female | 2850 | 1 | 6 | 0 | 0 |
| s41 | Non-harem | Male | 850 | 2 | 0 | 3 | 0 |
| s42 | Non-harem | Male | 3300 | 2 | 0 | 3 | 0 |
| s43 | Harem | Female | 3140 | 1 | 5 | 0 | 0 |
| s45 | Non-harem | Male | 815 | 2 | 0 | 3 | 0 |
| s46 | Non-harem | Female | 760 | 2 | 0 | 3 | 0 |
| s48 | Harem | Male | 2900 | 1 | 6 | 0 | 0 |

#### Supplementary Materials

Jose *et al.* Gut microbial community in proboscis monkeys (*Nasalis larvatus*): implications for effects of geographical and social factors

| | | | | | | | |
|---|---|---|---|---|---|---|---|
| s49 | Non-harem | Male | 2900 | 1 | 0 | 3 | 0 |
| s50 | Harem | Female | 2850 | 1 | 5 | 0 | 0 |
| s51 | Harem | Female | 2550 | 1 | 6 | 0 | 0 |
| s52 | Harem | Male | 1750 | 1 | 7 | 0 | 0 |
| s53 | Non-harem | Male | 3070 | 2 | 0 | 3 | 0 |
| s54 | Harem | Male | 2710 | 1 | 5 | 0 | 0 |
| s55 | Harem | Female | 2040 | 1 | 6 | 0 | 0 |
| s56 | Harem | Female | 2930 | 1 | 6 | 0 | 0 |
| s57 | Harem | Male | 2800 | 1 | 5 | 0 | 0 |
| s58 | Harem | Female | 2000 | 1 | 6 | 0 | 0 |
| s59 | Harem | Female | 1940 | 1 | 6 | 0 | 0 |
| s60 | Non-harem | Male | 1950 | 2 | 0 | 3 | 0 |
| s61 | Harem | Female | 300 | 1 | 5 | 0 | 0 |
| s62 | Harem | Male | 3100 | 1 | 6 | 0 | 0 |
| s63 | Harem | Female | 3100 | 1 | 6 | 0 | 0 |
| s64 | Harem | Female | 2000 | 1 | 5 | 0 | 0 |
| s65 | Harem | Male | 2000 | 1 | 5 | 0 | 0 |
| s66 | Harem | Female | 2870 | 1 | 5 | 0 | 0 |
| s67 | Harem | Female | 2405 | 1 | 7 | 0 | 0 |
| s68 | Non-harem | Female | 1400 | 2 | 0 | 3 | 0 |
| s69 | Harem | Male | 3000 | 1 | 5 | 0 | 0 |
| s70 | Non-harem | Male | 2310 | 2 | 0 | 3 | 0 |
| s71 | Non-harem | Female | 755 | 2 | 0 | 4 | 0 |
| s72 | Non-harem | Female | 2400 | 2 | 0 | 3 | 0 |
| s73 | Non-harem | Female | 2400 | 2 | 0 | 3 | 0 |
| s74 | Non-harem | Female | 2400 | 2 | 0 | 3 | 0 |
| s75 | Non-harem | Female | 300 | 2 | 0 | 3 | 0 |
| s77 | Non-harem | Male | 2720 | 2 | 0 | 3 | 0 |
| s78 | Harem | Female | 2640 | 1 | 5 | 0 | 0 |
| s79 | Harem | Male | 2310 | 1 | 5 | 0 | 0 |
| s80 | Harem | Female | 2000 | 1 | 6 | 0 | 0 |
| s81 | Harem | Female | 560 | 1 | 5 | 0 | 0 |
| s82 | Non-harem | Female | 3550 | 2 | 0 | 3 | 0 |
| s83 | Non-harem | Female | 3550 | 2 | 0 | 3 | 0 |
| s84 | Harem | Female | 3450 | 1 | 5 | 0 | 0 |
| s86 | Harem | Female | 2155 | 1 | 6 | 0 | 0 |
| s87 | Non-harem | Female | 2150 | 2 | 0 | 3 | 0 |
| s88 | Non-harem | Female | 2150 | 2 | 0 | 3 | 0 |
| s90 | Harem | Female | 560 | 1 | 6 | 0 | 0 |
| s91 | Harem | Female | 560 | 1 | 6 | 0 | 0 |
| s92 | Harem | Female | 560 | 1 | 6 | 0 | 0 |
| s93 | Harem | Female | 560 | 1 | 6 | 0 | 0 |

#### Supplementary Materials

Jose *et al.* Gut microbial community in proboscis monkeys (*Nasalis larvatus*): implications for effects of geographical and social factors

| | | | | | | | |
|---|---|---|---|---|---|---|---|
| s95 | Harem | Female | 560 | 1 | 6 | 0 | 0 |
| s96 | Harem | Male | 3890 | 1 | 6 | 0 | 0 |
| s97 | Harem | Male | 3740 | 1 | 4 | 0 | 0 |
| s98 | Non-harem | Female | 3450 | 2 | 0 | 2 | 0 |
| s99 | Harem | Male | 2850 | 1 | 6 | 0 | 0 |
| s100 | Non-harem | Male | 2800 | 2 | 0 | 3 | 0 |
| s101 | Non-harem | Female | 745 | 2 | 0 | 3 | 0 |
| s103 | Harem | Female | 3460 | 1 | 6 | 0 | 0 |
| s105 | Harem | Female | 2920 | 1 | 4 | 0 | 0 |
| s106 | Harem | Female | 2920 | 1 | 4 | 0 | 0 |
| s107 | Harem | Female | 3530 | 1 | 6 | 0 | 0 |
| s108 | Harem | Female | 3530 | 1 | 6 | 0 | 0 |
| s111 | Non-harem | Male | 900 | 2 | 0 | 3 | 0 |
| s114 | Non-harem | Female | 2270 | 2 | 0 | 3 | 0 |
| s115 | Non-harem | Male | 2270 | 2 | 0 | 3 | 0 |
| s116 | Harem | Female | 1950 | 1 | 6 | 0 | 0 |
| s117 | Harem | Female | 1950 | 1 | 6 | 0 | 0 |
| s118 | Harem | Female | 1950 | 1 | 6 | 0 | 0 |
| s119 | Harem | Female | 200 | 1 | 5 | 0 | 0 |
| s120 | Harem | Female | 200 | 1 | 5 | 0 | 0 |
| s121 | Harem | Female | 3950 | 1 | 5 | 0 | 0 |
| s122 | Harem | Female | 3950 | 1 | 5 | 0 | 0 |
| s124 | Non-harem | Male | 3700 | 2 | 0 | 3 | 0 |
| s125 | Non-harem | Female | 3450 | 2 | 0 | 2 | 0 |
| s126 | Harem | Female | 4320 | 1 | 6 | 0 | 0 |
| s127 | Non-harem | Female | 600 | 2 | 0 | 3 | 0 |
| s128 | Non-harem | Female | 600 | 2 | 0 | 3 | 0 |
| s129 | Harem | Female | 580 | 1 | 5 | 0 | 0 |
| s130 | Harem | Female | 580 | 1 | 5 | 0 | 0 |
| s131 | Harem | Female | 4150 | 1 | 6 | 0 | 0 |
| s132 | Harem | Female | 2000 | 1 | 5 | 0 | 0 |
| s133 | Harem | Female | 1450 | 1 | 5 | 0 | 0 |
| s135 | Non-harem | Female | 550 | 2 | 0 | 3 | 0 |
| s136 | Harem | Male | 3810 | 1 | 6 | 0 | 0 |
| s137 | Harem | Female | 3760 | 1 | 5 | 0 | 0 |
| s138 | Harem | Female | 3630 | 1 | 6 | 0 | 0 |
| s139 | Non-harem | Male | 3300 | 2 | 0 | 3 | 0 |
| s140 | Harem | Female | 2800 | 1 | 5 | 0 | 0 |
| s141 | Harem | Male | 990 | 1 | 6 | 0 | 0 |
| s142 | Non-harem | Male | 3300 | 2 | 0 | 3 | 0 |
| s143 | Harem | Female | 2890 | 1 | 7 | 0 | 0 |
| s144 | Harem | Female | 2800 | 1 | 4 | 0 | 0 |
| s145 | Non-harem | Male | 2440 | 1 | 0 | 0 | 0 |

#### Supplementary Materials

Jose *et al.* Gut microbial community in proboscis monkeys (*Nasalis larvatus*): implications for effects of geographical and social factors

| | | | | | | | |
|---|---|---|---|---|---|---|---|
| s147 | Harem | Female | 610 | 1 | 7 | 0 | 0 |
| s148 | Harem | Female | 200 | 1 | 6 | 0 | 0 |
| s149 | Non-harem | Female | 1700 | 2 | 0 | 3 | 0 |
| s150 | Non-harem | Female | 1700 | 2 | 0 | 3 | 0 |
| s151 | Harem | Female | 1660 | 1 | 6 | 0 | 0 |
| s152 | Harem | Female | 1660 | 1 | 6 | 0 | 0 |
| s153 | Harem | Female | 1660 | 1 | 6 | 0 | 0 |
| s154 | Harem | Female | 1660 | 1 | 6 | 0 | 0 |
| s155 | Harem | Female | 1660 | 1 | 6 | 0 | 0 |
| s156 | Harem | Female | 1660 | 1 | 6 | 0 | 0 |
| s157 | Harem | Female | 1660 | 1 | 6 | 0 | 0 |
| s158 | Harem | Female | 1660 | 1 | 6 | 0 | 0 |
| s159 | Harem | Male | 1660 | 1 | 6 | 0 | 0 |
| s160 | Harem | Female | 2090 | 1 | 6 | 0 | 0 |
| s161 | Harem | Female | 2090 | 1 | 6 | 0 | 0 |
| s162 | Harem | Female | 2090 | 1 | 6 | 0 | 0 |
| s163 | Harem | Female | 2090 | 1 | 6 | 0 | 0 |
| s164 | Harem | Female | 2090 | 1 | 6 | 0 | 0 |
| s165 | Harem | Female | 1700 | 1 | 8 | 0 | 0 |
| s167 | Harem | Female | 1700 | 1 | 8 | 0 | 0 |
| s168 | Harem | Female | 1700 | 1 | 8 | 0 | 0 |
| s169 | Harem | Female | 1700 | 1 | 8 | 0 | 0 |
| s170 | Harem | Female | 1700 | 1 | 8 | 0 | 0 |
| s171 | Harem | Female | 1700 | 1 | 8 | 0 | 0 |
| s172 | Harem | Female | 1700 | 1 | 8 | 0 | 0 |
| s173 | Harem | Female | 1700 | 1 | 8 | 0 | 0 |
| s174 | Harem | Female | 1700 | 1 | 8 | 0 | 0 |
| s175 | Harem | Female | 1700 | 1 | 8 | 0 | 0 |
| s176 | Harem | Male | 2400 | 1 | 6 | 0 | 0 |
| s177 | Harem | Male | 2400 | 1 | 6 | 0 | 0 |
| s178 | Harem | Female | 2400 | 1 | 6 | 0 | 0 |
| s179 | Harem | Male | 2400 | 1 | 6 | 0 | 0 |
| s180 | Harem | Female | 2400 | 1 | 6 | 0 | 0 |
| s183 | Harem | Female | 2400 | 1 | 6 | 0 | 0 |
| s184 | Harem | Female | 1945 | 1 | 8 | 0 | 0 |
| s185 | Harem | Female | 1945 | 1 | 8 | 0 | 0 |
| s186 | Harem | Female | 1945 | 1 | 8 | 0 | 0 |
| s187 | Non-harem | Male | 850 | 2 | 0 | 2 | 0 |
| s188 | Non-harem | Male | 850 | 2 | 0 | 2 | 0 |
| s189 | Non-harem | Male | 850 | 2 | 0 | 2 | 0 |
| s190 | Non-harem | Male | 850 | 2 | 0 | 2 | 0 |
| s192 | Non-harem | Male | 300 | 2 | 0 | 3 | 0 |
| s193 | Non-harem | Male | 300 | 2 | 0 | 3 | 0 |
| s194 | Non-harem | Male | 300 | 2 | 0 | 3 | 0 |
| s196 | Non-harem | Male | 300 | 2 | 0 | 3 | 0 |
| s197 | Non-harem | Male | 300 | 2 | 0 | 3 | 0 |

| | | | | | | | |
|---|---|---|---|---|---|---|---|
| s198 | Non-harem | Male | 300 | 2 | 0 | 3 | 0 |
| s199 | Non-harem | Male | 300 | 2 | 0 | 3 | 0 |
| s200 | Non-harem | Female | 300 | 2 | 0 | 3 | 0 |
| s201 | Non-harem | Male | 300 | 2 | 0 | 3 | 0 |
| s202 | Harem | Female | 2100 | 1 | 8 | 0 | 0 |
| s203 | Harem | Female | 2100 | 1 | 8 | 0 | 0 |
| s204 | Harem | Female | 2100 | 1 | 8 | 0 | 0 |
| s205 | Harem | Female | 2100 | 1 | 8 | 0 | 0 |
| s206 | Harem | Female | 2100 | 1 | 8 | 0 | 0 |
| s207 | Harem | Male | 2100 | 1 | 8 | 0 | 0 |
| s208 | Harem | Female | 2100 | 1 | 8 | 0 | 0 |
| s209 | Harem | Male | 2100 | 1 | 8 | 0 | 0 |
| s210 | Non-harem | Female | 600 | 2 | 0 | 4 | 0 |
| s211 | Non-harem | Male | 600 | 2 | 0 | 4 | 0 |
| s213 | Non-harem | Male | 600 | 2 | 0 | 4 | 0 |
| s214 | Non-harem | Male | 600 | 2 | 0 | 4 | 0 |
| s215 | Non-harem | Female | 400 | 2 | 0 | 3 | 0 |
| s216 | Non-harem | Female | 400 | 2 | 0 | 3 | 0 |
| s217 | Non-harem | Female | 400 | 2 | 0 | 3 | 0 |
| s218 | Non-harem | Male | 400 | 2 | 0 | 3 | 0 |
| s219 | Non-harem | Female | 400 | 2 | 0 | 3 | 0 |
| s220 | Non-harem | Female | 400 | 2 | 0 | 3 | 0 |
| s222 | Non-harem | Female | 400 | 2 | 0 | 3 | 0 |
| s223 | Non-harem | Female | 400 | 2 | 0 | 3 | 0 |
| s224 | Non-harem | Female | 400 | 2 | 0 | 3 | 0 |
| s225 | Non-harem | Female | 400 | 2 | 0 | 3 | 0 |
| s226 | Non-harem | Male | 400 | 2 | 0 | 3 | 0 |
| s227 | Harem | Female | 3400 | 1 | 7 | 0 | 0 |
| s228 | Harem | Female | 3400 | 1 | 7 | 0 | 0 |
| s229 | Harem | Female | 3400 | 1 | 7 | 0 | 0 |
| s230 | Harem | Female | 3400 | 1 | 7 | 0 | 0 |
| s232 | Harem | Female | 3400 | 1 | 7 | 0 | 0 |
| s233 | Harem | Female | 3400 | 1 | 7 | 0 | 0 |
| s234 | Harem | Female | 3400 | 1 | 7 | 0 | 0 |
| s235 | Harem | Female | 2100 | 1 | 6 | 0 | 0 |
| s236 | Harem | Female | 2100 | 1 | 6 | 0 | 0 |
| s237 | Harem | Female | 2100 | 1 | 6 | 0 | 0 |
| s238 | Harem | Male | 2100 | 1 | 6 | 0 | 0 |

#### Supplementary Materials

Jose *et al.* Gut microbial community in proboscis monkeys (*Nasalis larvatus*): implications for effects of geographical and social factors

| | | | | | | | |
|---|---|---|---|---|---|---|---|
| s239 | Harem | Female | 2100 | 1 | 6 | 0 | 0 |
| s240 | Harem | Female | 2750 | 1 | 6 | 0 | 0 |
| s241 | Harem | Female | 2750 | 1 | 6 | 0 | 0 |
| s243 | Harem | Female | 2750 | 1 | 6 | 0 | 0 |
| s244 | Harem | Female | 2750 | 1 | 6 | 0 | 0 |
| s245 | Harem | Female | 2750 | 1 | 6 | 0 | 0 |
| s247 | Harem | Female | 2450 | 1 | 9 | 0 | 0 |
| s248 | Harem | Male | 2450 | 1 | 9 | 0 | 0 |
| s249 | Harem | Female | 2450 | 1 | 9 | 0 | 0 |
| s250 | Harem | Female | 2450 | 1 | 9 | 0 | 0 |
| s252 | Harem | Female | 2450 | 1 | 9 | 0 | 0 |
| s255 | Non-harem | Male | 550 | 2 | 0 | 2 | 0 |
| s256 | Non-harem | Female | 550 | 2 | 0 | 2 | 0 |
| s258 | Non-harem | Female | 550 | 2 | 0 | 2 | 0 |
| s259 | Non-harem | Female | 550 | 2 | 0 | 2 | 0 |
| s260 | Harem | Female | 2450 | 1 | 4 | 0 | 0 |
| s261 | Harem | Male | 2450 | 1 | 4 | 0 | 0 |
| s262 | Harem | Male | 2450 | 1 | 4 | 0 | 0 |
| s263 | Harem | Female | 2450 | 1 | 4 | 0 | 0 |
| s264 | Harem | Female | 2450 | 1 | 4 | 0 | 0 |
| s265 | Harem | Female | 200 | 1 | 6 | 0 | 0 |
| s266 | Harem | Female | 200 | 1 | 6 | 0 | 0 |
| s267 | Harem | Female | 200 | 1 | 6 | 0 | 0 |
| s268 | Harem | Female | 1860 | 1 | 10 | 0 | 0 |
| s269 | Harem | Female | 200 | 1 | 6 | 0 | 0 |
| s271 | Harem | Female | 200 | 1 | 6 | 0 | 0 |
| s272 | Harem | Female | 200 | 1 | 6 | 0 | 0 |
| s273 | Harem | Female | 200 | 1 | 6 | 0 | 0 |
| s274 | Harem | Female | 200 | 1 | 6 | 0 | 0 |
| s275 | Non-harem | Female | 190 | 2 | 0 | 4 | 0 |
| s276 | Non-harem | Female | 190 | 2 | 0 | 4 | 0 |
| s277 | Non-harem | Male | 190 | 2 | 0 | 4 | 0 |
| s278 | Non-harem | Male | 200 | 2 | 0 | 3 | 0 |
| s279 | Non-harem | Male | 200 | 2 | 0 | 3 | 0 |
| s280 | Non-harem | Male | 200 | 2 | 0 | 3 | 0 |
| s281 | Non-harem | Female | 200 | 2 | 0 | 3 | 0 |
| s285 | Non-harem | Male | 200 | 2 | 0 | 3 | 0 |
| s287 | Non-harem | Male | 200 | 2 | 0 | 3 | 0 |
| s288 | Non-harem | Female | 910 | 2 | 0 | 3 | 0 |
| s289 | Non-harem | Male | 910 | 2 | 0 | 3 | 0 |
| s290 | Non-harem | Female | 910 | 2 | 0 | 3 | 0 |

#### Supplementary Materials

Jose *et al.* Gut microbial community in proboscis monkeys (*Nasalis larvatus*): implications for effects of geographical and social factors

| | | | | | | | |
|---|---|---|---|---|---|---|---|
| s291 | Non-harem | Female | 910 | 2 | 0 | 3 | 0 |
| s292 | Non-harem | Male | 910 | 2 | 0 | 3 | 0 |
| s293 | Non-harem | Female | 910 | 2 | 0 | 3 | 0 |
| s294 | Non-harem | Male | 910 | 2 | 0 | 3 | 0 |
| s295 | Harem | Male | 2760 | 1 | 5 | 0 | 0 |
| s296 | Harem | Male | 2760 | 1 | 5 | 0 | 0 |
| s297 | Harem | Male | 2760 | 1 | 5 | 0 | 0 |
| s298 | Harem | Male | 2760 | 1 | 5 | 0 | 0 |
| s299 | Harem | Female | 2010 | 1 | 4 | 0 | 0 |
| s300 | Harem | Female | 2010 | 1 | 4 | 0 | 0 |
| s301 | Harem | Female | 2010 | 1 | 4 | 0 | 0 |
| s302 | Harem | Female | 2010 | 1 | 4 | 0 | 0 |
| s303 | Harem | Female | 2010 | 1 | 4 | 0 | 0 |
| s304 | Harem | Female | 2010 | 1 | 4 | 0 | 0 |
| s305 | Harem | Female | 2010 | 1 | 4 | 0 | 0 |
| s306 | Harem | Female | 2010 | 1 | 4 | 0 | 0 |

[*]Although there was only one adult male in the group, as five immature males (juveniles) were observed in the group, we considered the group type as a non-harem group.

### Supplementary Materials

Jose *et al.* Gut microbial community in proboscis monkeys (*Nasalis larvatus*): implications for effects of geographical and social factors

#### Supplementary Material 9: Vegetation survey and plant diversity

In 2005, to facilitate the observation and tracking of primates in the riverine forest, trails measuring 200–500 m in length and 1 m in width were established at 500 m intervals on both sides of the Menanggul River by clearing the forest floor undergrowth (Supplementary Material 1). Trees with a diameter at breast height (DBH) of ≥ 10 cm and vines with a diameter of ≥ 5 cm located either on the trail or within 1 m from the trail edge were identified and marked (a sample area width of 3 m). Taxonomic identification of all labeled trees and vines was conducted with the assistance of the Forest Research Center, Sandakan, Sabah. A total of 16 trails (TR1-16) were established, all of which were 500 m in length, except for TR 2 (200 m long), TR 8 (250 m long), and TR 4, TR 6, and TR 10 (400 m long) [1]. The cumulative survey area encompassed 2.15 ha (i.e., 7150 m × 3 m). Along the 16 trails, we marked 1,645 trees and 497 vines, representing 180 species, 124 genera, and 46 families. Among the 2,142 trees and vines identified along the trails, 1,902 were potential food sources for the proboscis monkeys [2].

Shannon's diversity index (*H'*) was computed from the total count of each plant species along survey trails, yielding a mean value of 3.45 (SD = 0.26) across all trails. Based on our previous report [3], we compared the plant diversity of TR1-TR8 ( trails from the mouth of the river to 2,000 m upstream) and TR9-TR16 (trails from 2,000 m to 4,000 further upstream), as there are two different proboscis monkey communities between those area. A comparison of the mean Shannon's diversity index between the downstream area including eight trails (TR1-TR8) and upstream area including eight trails (TR9-TR16) revealed a trend towards lower diversity indices in the downstream area, although this disparity did not achieve statistical significance ($U= 23$; $p=0.38$), i.e. downstream: $H'=3.37$ (SD=0.29), upstream: $H'=3.53$ (SD=0.22). It is noteworthy that the trail established on the north side of the river near the river mouth (TR2), where deforestation by oil palm plantations is most evident, had the lowest diversity index of 2.80. Shannon's diversity index computed based on potential food plants for proboscis monkeys exhibited a similar trend: the overall mean was 3.32 (SD = 0.29), with downstream areas showing a mean of 3.22 (SD = 0.35) and upstream areas showing a mean of 3.41 (SD = 0.18). A comparison of the mean diversity index between downstream and upstream areas also did not reveal a significant difference ($U = 20$; $p = 0.23$). Additionally, TR2 exhibited the lowest potential food plant diversity index of 2.50.
